# The structure and spontaneous curvature of clathrin lattices at the plasma membrane

**DOI:** 10.1101/2020.07.18.207258

**Authors:** Kem A. Sochacki, Bridgette L. Heine, Gideon J. Haber, John R. Jimah, Bijeta Prasai, Marco A. Alfonzo-Mendez, Aleah D. Roberts, Agila Somasundaram, Jenny E. Hinshaw, Justin W. Taraska

## Abstract

Clathrin mediated endocytosis is the primary pathway for receptor and cargo internalization in eukaryotic cells. It is characterized by a polyhedral clathrin lattice that coats budding membranes. The mechanism and control of lattice assembly, curvature, and vesicle formation at the plasma membrane has been a matter of longstanding debate. Here, we use platinum replica and cryo electron microscopy and tomography to present a global structural framework of the pathway. We determine the shape and size parameters common to clathrin-mediated endocytosis. We show that clathrin sites maintain a constant surface area during curvature across multiple cell lines. Flat clathrin is present in all cells and spontaneously curves into coated pits without additional energy sources or recruited factors. Finally, we attribute curvature generation to loosely connected and pentagon-containing flat lattices that can rapidly curve when a flattening force is released. Together, these data present a new universal mechanistic model of clathrin-mediated endocytosis.

## Introduction

A key challenge for eukaryotic cells is the need to collect specific extracellular or integral membrane materials including nutrients, hormones, membrane-bound receptors, and proteins, and selectively incorporate them into small vesicles for transport into the cell. In eukaryotic cells, this task is largely accomplished by a process called clathrin mediated endocytosis (CME). Clathrin, the pathway’s protein namesake, is a cytosolic hetero-hexamer made of three heavy and three light chains that assembles as a three-legged pinwheel or “triskelion”. Triskelia further join into an interwoven honeycomb-like lattice made mostly of hexagons and pentagons. Along with dozens of other adaptors and regulatory proteins, clathrin lattices form a dynamic membrane-bound coat. Many of these factors, and their biophysical features, are thought to collectively induce or stabilize membrane curvature.^1-5^ As a result, clathrin coats and bends the membrane into a nanoscale cargo-filled sphere. Once inside the cell, clathrin is released and the vesicle is freed from its clathrin cage.

Currently, there are two competing structural models for how clathrin lattices assemble and mature. These are 1) the constant curvature model and 2) the constant area model. In the first, clathrin lattices grow like a rising sun with a fixed radius of curvature.^6^ Thus, flat clathrin lattices either do not exist or are not capable of endocytosis. In the second, clathrin lattices grow flat to their terminal surface area and then bend to create a vesicle.^7^ Mixtures of these models have been observed recently; clathrin sites growing flat, growing curved, or switching between behaviors, even within the same cell.^8, 9^ Indeed, clathrin imaging in live cells exhibits varied lifetime dynamics, consistent with this mixed-model of endocytosis.^10, 11^ To test these models and determine their universality, a large-scale comparison of clathrin structure within and among cell lines is needed. However, these data have been missing—their absence likely due to the challenges in acquiring, segmenting, and analyzing large amounts of EM data at the nanoscale across many cells and types of cells.

While the two major models of CME are still openly debated, recent correlative and live-cell fluorescence microscopies have supported the idea that clathrin can grow flat and then curve in some cells.^8, 9, 12^ This is surprising considering the atomic structure of clathrin baskets observed from cryo electron microscopy (cryoEM).^13, 14^ These minimal polyhedral baskets are characterized by a high degree of interdigitation with each lattice hub incorporating portions of seven different triskelia. The result is that even though lone triskelia are quite flexible, the baskets are structured with enough rigidity to perform cryoEM averaging with better than 10 Angstroms of consensus.^13-15^ Conversely, cryoEM data on flat clathrin structures is lacking. Combining lower resolution EM images and an extrapolation from cryoEM baskets, flat clathrin is generally modeled as a tightly connected, rigid grid of hexagons.^7, 13, 16^ Yet, according to Euler’s theorem, creating a sphere from a flat hexagonal lattice requires the insertion of 12 pentagons.^17^ This would impose substantial structural and energetic barriers to curvature.^16, 18^ To overcome these barriers, the dominant model for flat clathrin lattice bending involves a controlled energy-dependent exchange of clathrin subunits. This would permit the lattice rearrangements necessary for curvature.^12^

Adjacent to its role in endocytosis, flat clathrin lattices have also been proposed to play other important physiological roles. For example, flat clathrin can act as an integrin-containing adhesion site,^19^ connecting retraction fibers to the extracellular substrate during mitosis,^20^ adhering to engulfed particles during phagocytosis,^21^ gripping contractile fibers in skeletal muscle,^22, 23^ and aiding cell motility.^24-26^ Furthermore, an increase of flat clathrin during signaling has suggested that it could act as a dynamic signaling platform.^27^ Thus, understanding the full structural landscape of clathrin —from flat to curved— is important for understanding not only endocytosis, but also clathrin’s expanding role in the cell.

Here, we study the universal structural features of clathrin lattices in mammalian cells. We examine the size, shape, and density of clathrin at the plasma membrane of eight commonly used mammalian cell types. We find that all cells contain mixtures of flat, domed, and spherical patches of clathrin. We show that lattices bend while maintaining a relatively constant surface area during curvature. We observe that flat clathrin spontaneously curves when the top of the cell and the cytoplasm are removed. Thus, added energy and proteins are not necessary for lattice curvature. We count pentagons within flat and domed structures and find that flat clathrin contains many pentagons—minimizing the need for gross lattice reorganization during curvature. Finally, we examine molecular-scale interactions of triskelia in individual flat lattices using cryo-electron tomography (cryoET). We observe that flat clathrin is loosely assembled with many structural defects. We propose that flat clathrin is loaded like a spring or locked Brownian ratchet, harboring potential energy in the form of a strained or loosely connected network of inter-triskelion interactions. Collectively, these data define a universal structural and regulatory framework for clathrin lattice assembly, growth, and curvature.

## Results

### Quantitative structural assessment of clathrin in HeLa cells

Flat clathrin bends into endocytic vesicles in both human skin melanoma (SK-MEL-2) and green monkey kidney fibroblasts (BS-C-1).^8, 9, 12^ Yet, these cell lines have been noted for a lack of large flat clathrin plaques.^9, 28^ HeLa cells, however, contain many large flat clathrin plaques.^28, 29^ To study the effect of these structures on the stages of endocytosis, we analyzed the global structure and distribution of clathrin across the plasma membrane in HeLa cells. To do this, we used platinum replica electron microscopy (PREM) which provides a uniquely high contrast, high resolution, wide-scale view of the inner surface of the plasma membrane.^7, 30^ Cells grown on coated glass coverslips were disrupted with a sheer force in fixative to stabilize, isolate, coat, and then image the adherent membrane (Fig. 1a).^31, 32^ From these PREM images every visible clathrin structure could be classified as flat (no noticeable curvature), domed (curved less than a hemisphere so the edge is still visible from above), or spherical (curved beyond a hemisphere so the edge is no longer visible from above) (Fig. 1b-d).

**Figure 1.**
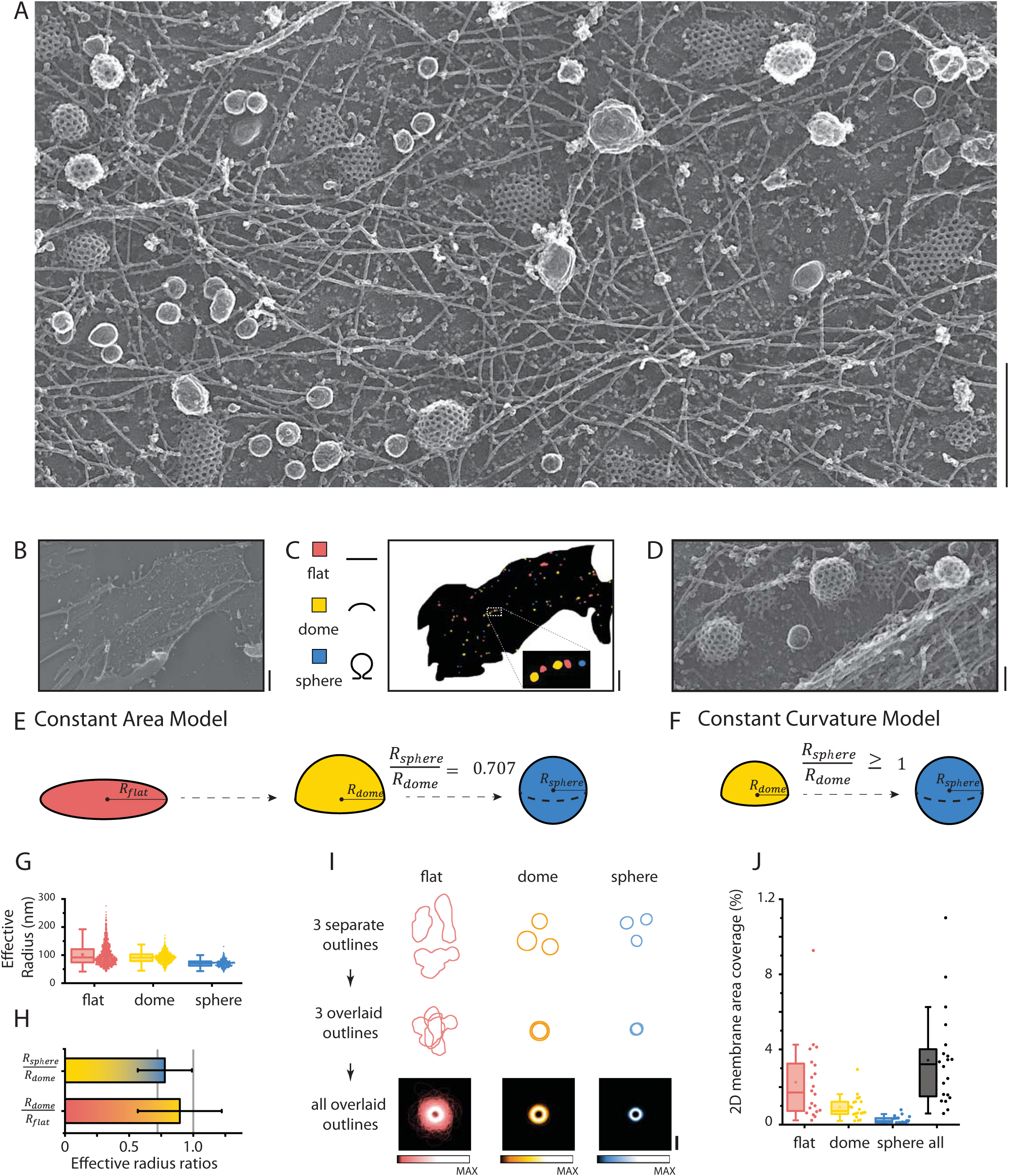
HeLa cells exhibit constant area between dome and sphere. (A) A representative crop from a PREM image of an unroofed HeLa cell. (B) A montaged area of a full membrane is segmented to create a mask as in (C) with total membrane area (non-white), flat (pink), domed (yellow), and sphere (blue) structures. The zoomed-in break-out box shows all three clathrin classifications. The image of these structures is shown in (D). (E) The constant area model states that clathrin grows in the flat state. The surface area of clathrin remains constant as the structure curves into a vesicle. (F) The constant curvature model states that clathrin structures grow with a constant radius of curvature. In this case, endocyticly active clathrin is never flat. The two models can be differentiated by the relationship between the radii of domed (R_dome_) and sphere (R_sphere_) structures with the ratios shown. (G) Effective radius (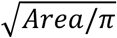 where *Area* is 2D projected area) of each structure and box plots are shown for HeLa cells. (H) Radius ratios are shown as a bar plot (mean ± standard deviation). The gray lines show the expected relationships for 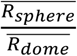 for the constant area model (0.707) or constant curvature model (≥1) respectively. (I) Clathrin structure outlines are combined to summarize size and shape differences. To illustrate how this was done, we show that the outlines are collected, centered, and summed to create the final summary. The color scale for intensity is shown under the images with max intensity labelled. (J) Percentage membrane area coverage is shown for each type of clathrin structure and the sum of all clathrin structures. N_flat_=1197, N_dome_=666, N_sphere_=322; N_cells_ = 20. For box plots, box is interquartile range, center line is median, center circle is mean, whiskers include all data excluding outliers. Scale bars in A, B, C, D, I are 500 nm, 2 µm, 2 µm, 100 nm, 200 nm respectively.

Next, to determine how clathrin curves we used the fact that the geometric relationship between a dome and sphere radius (*R*_*dome*_ and *R*_*sphere*_) differs between the two proposed models of endocytosis. Specifically, in the constant area model, a hemisphere that curves into a sphere while maintaining a constant surface area will have a larger radius by a factor of 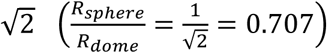 (Fig. 1e). In the constant curvature model, however, where surface area increases during growth, the ratio of radii between a dome and sphere is always greater than or equal to one 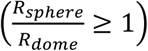 (Fig. 1f). In Figure 1g we measured the projected image areas of thousands of clathrin structures and converted the areas to effective radii 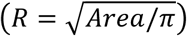. This conversion calculates radius by estimating the segmented object as circular. Figure 1h shows that from these values, in HeLa cells, the sphere to dome ratio is far less than one 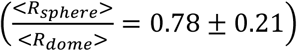 (mean ± st. dev) (Fig. 1h). These data are consistent with the constant area model and do not support the constant curvature model where many domes would be small and cannot exceed the size of a vesicle.

We next tested the step of flat to dome transition. A flat disk-shaped clathrin site that transitions into a dome without a change in surface area would result in the same relationship between radii as discussed 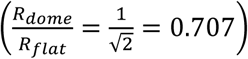. However, this cannot be used to differentiate between the two opposing models. Specifically, the constant curvature model predicts that 1) flat structures do not exist or 2) they are not endocytic. The constant area model proposes that growth during the flat stage would result in small flat intermediates before curvature. In HeLa cells, we found the measured dome to flat radii ratio was larger than 0.71 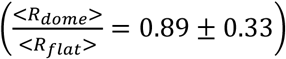 (Fig. 1h). Therefore, dome structures have a slightly larger surface area than flat structures on average. Thus, small flat structures are present in these membranes consistent with progressive growth at the flat stage. Figure 1i shows the segmentation outlines of all clathrin structures to provide a visual summary of the two-dimensional projection area observed. Of note, for domes and spheres, surface area is 2x and 4x the two-dimensional projection area respectively. Flat clathrin is clearly more heterogenous in size and shape as expected for random lattice growth. We also measured the percentage of total membrane area occupied by clathrin objects (Fig. 1j). Flat clathrin structures occupy a much larger percentage of membrane area than domes or spheres (2 +/- 2%, 0.9 +/− 0.6%, 0.3+/−0.2% respectively). Therefore, even though flat clathrin has a smaller surface area on average than domes, as a population they occupy more two-dimensional space on HeLa cell membranes than domes and spheres combined. Thus, in HeLa cells, flat clathrin is a prominent feature of the plasma membrane.

### Density of flat clathrin structures is highly variable among cell lines

To probe the significance of these structural relationships, we determined which features varied among several different cell lines. We chose seven commonly used cell lines that covered multiple species and cell types: 1) 3T3-L1 mouse embryo fibroblast cells, 2) BS-C-1 green monkey kidney cells, 3) L6 rat myoblasts, 4) MCF7 human mammary gland adenocarcinoma cells, 5) PC12-GR5 rat adrenal gland cells, 6) RAW264.7 mouse monocyte/macrophages, and 7) U-87 MG human brain glioblastoma cells. In PREM images of these cells, clathrin was segmented as before (Fig. 2a-b). The reproducibility of manual segmentation was tested and confirmed within 9%, 5%, 3% error for flat, dome, and sphere radius measurements respectively, and 10%, 15% and 26% for flat, dome, and sphere, membrane area percentage measurements respectively (Fig. S1). Flat, domed, and spherical clathrin structures were visible in all cells. Thus, flat structures are not unique to rare or specific cell types. Representative segmentation masks illustrate the large diversity in densities (Fig. 2b). In box plots of total clathrin area occupation for flat, dome, and sphere subclasses, there is no clear trend among cell lines (Fig. 2c-e, Fig. S2a-b). Spheres occupy between 0.11% (L6) to 0.62% (PC-12) (mean) membrane area, a 5.6-fold range. Dome structures occupy between 0.72% (3T3) and 2.3% (MCF7) (mean) membrane area, a 3.2-fold range. Notably, flat structures occupy between 0.51% (BS-C-1) to 11% (MCF7) (mean) of membrane area, a 22-fold difference. This variability is consistent with the diverse lifetimes of clathrin structures between cells reported with fluorescence microscopy and highlight the architectural flexibility of the clathrin system.^10, 11, 33^ A cell line frequently used to study CME by microscopy, BS-C-1, has the lowest total clathrin membrane area at 1.5±0.8%, 2-fold smaller than the next highest cell type, 3T3 cells, and 9-fold smaller than the highest occupied membranes in MCF7 cells (Fig. S2a). Thus, clathrin is varied in both shape and density across commonly used mammalian cell types.

**Figure 2.**
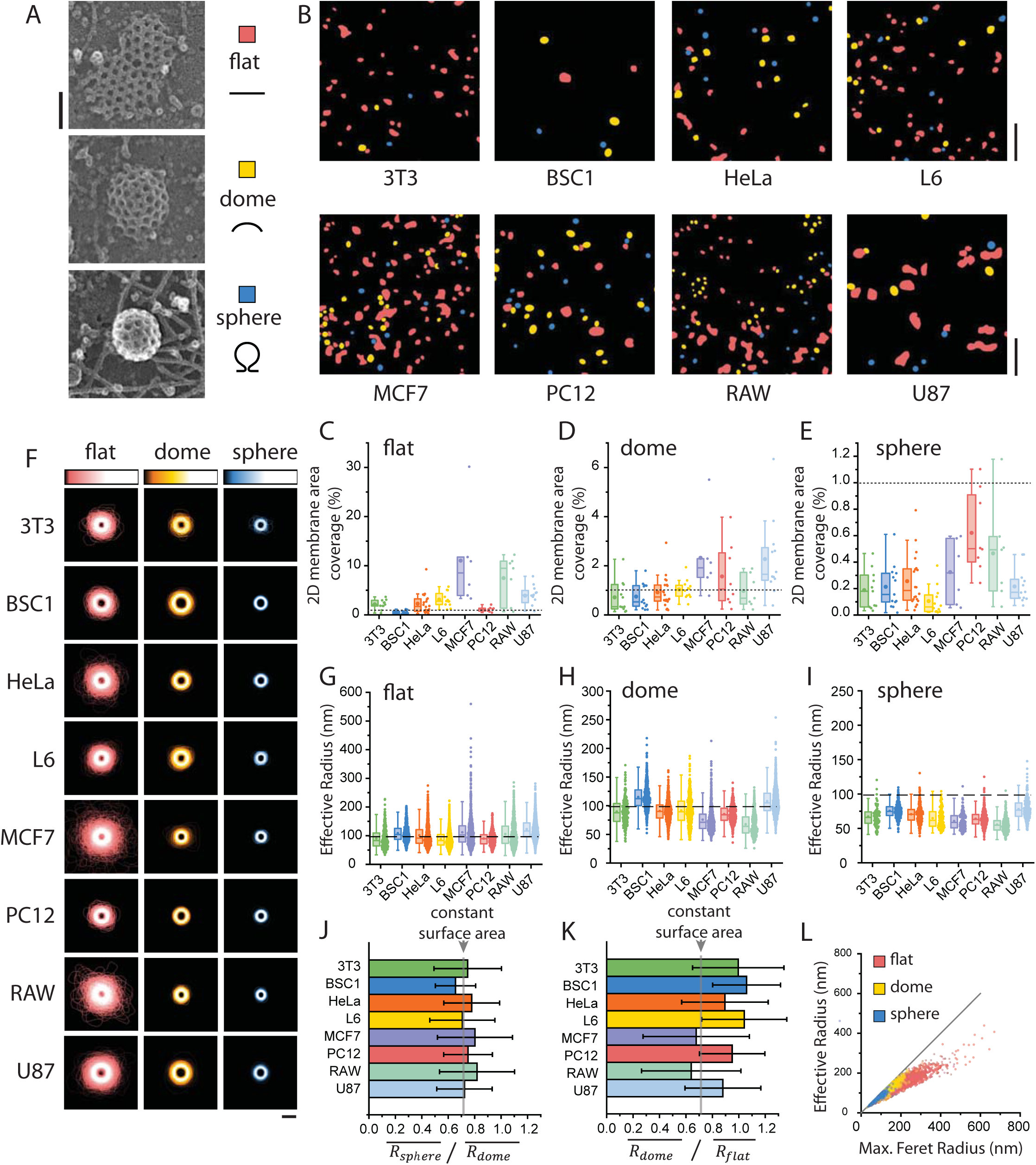
Eight different cell lines all exhibit constant area between dome and sphere but diversity in flat structures. (A) Examples of structures that would be segmented as flat, dome, or sphere. (B) Examples of segmentation in eight different cell lines. The membrane area fraction of (C) flat, (D) dome, and (E) sphere categories depicts the total area on each membrane occupied by that clathrin class divided by the total analyzed membrane area. The dotted line references 1% of the membrane area to show scale change. (F) Outline summaries for each cell line are shown as in Fig. 1i. The effective radius for (G) flat, (H) dome, and (I) sphere categories are shown. The dashed lines reference 100 nm to show scale change. (J) 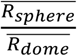 for each cell line is shown with average and standard deviation. (K) 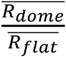 for each cell line is shown with average and standard deviation. The ^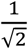^ relationship expected for constant surface area is referenced by a gray line in J-K. (L) Effective radius vs maximum Feret radius from all cell lines illustrates that flat structures are not circular. 3T3 N_flat_=597, N_dome_=211, N_sphere_=79, N_cells_ = 11; BSC1 N_flat_=568, N_dome_=694, N_sphere_=469, N_cells_ = 13; HeLa N_flat_=1197, N_dome_=666, N_sphere_=322, N_cells_ = 20; L6 N_flat_=2042, N_dome_=631, N_sphere_=141, N_cells_ = 12; MCF7 N_flat_=1471, N_dome_=705, N_sphere_=148, N_cells_ = 6; PC12 N_flat_=441, N_dome_=570, N_sphere_=474, N_cells_ = 8; RAW N_flat_=1139, N_dome_=454, N_sphere_=266, N_cells_ = 7; U87 N_flat_=1996, N_dome_=1329, N_sphere_=300, N_cells_ = 12. For box plots, box is interquartile range, center line is median, center circle is mean, whiskers include all data excluding outliers. Data shown to the right of box plots is from each separate cell in D-G and for each separate clathrin structure in I-K. Scale bars in A, B, and F are 100nm, 1 micron, and 200 nm respectively.

### Clathrin consistently bends from dome to sphere with a constant surface area across cell types

Figure 2f shows a trend amongst cell lines when the summed outlines of clathrin structures in all eight cell lines are measured. Clathrin spheres are consistently smaller than clathrin domes. Box plots of effective radii show that cell lines with larger spheres (U-87 and BS-C-1) also have larger domes. Cell lines with smaller spheres (MCF7 and RAW264.7) have smaller domes. The range in average sphere radius varies from 55.4 ± 11.1 nm (RAW264.7) to 78.3 ± 14.9 nm (U-87), effectively doubling the surface area. All cell types, however, exhibit a mean radius larger than the classic soccer ball shape (truncated icosohedron, 60 triskelia, 47 nm radius) or cryoEM cage structures (36, 37 triskelia, ∼35 nm radius).^13, 14^ Remarkably, the 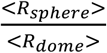 ratio is consistently near ^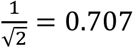^ (Fig. 2j, Fig. S2c) as it was for HeLa.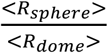 ranged from 0.66 ± 0.15 in BS-C-1 cells to 0.82 ± 0.28 in RAW264.7 cells. Thus, the structural relationship between domes and spheres in all cell lines fits with a model where clathrin maintains a near constant surface area during curvature.

Figure 2k shows that, in contrast, there is not a constant relationship between the size of flat structures and dome structures across cell types. The average radius of flat structures varies from 88.0 ±28.3 nm in L6 to 122.7 ± 37.4 nm in U-87 cells, a 2-fold change. Yet, having smaller flat structures does not correlate with smaller domes. Specifically, most cell lines (3T3-L1, BS-C-1, HeLa, L6, PC-12, U-87) exhibit a sphere to dome radius ratio greater than 0.707, indicating larger surface area on average in domes than flat structures. Two cell lines (MCF7, RAW264.7) exhibit ratios consistent with smaller surface area on average in domes versus flat structures. We also quantitatively assessed the overall shape of clathrin structures by comparing their effective radius versus their maximum Feret radius (half of the largest calipered distance in the structure) (Fig. 2l, Fig. S2d-f). For circular objects this ratio should be near one. For elongated objects this value will be less than one. For domes and spheres, these values support a circular morphology and fall along a line with the slope of one. However, flat structures fall distinctly below this line, emphasizing their irregular shapes. Therefore, 1) the size of flat structures does not relate to the sizes of domes or spheres, 2) they exhibit vast heterogeneity in density and shape, and 3) they tend to grow in non-spherical geometries.

### Spontaneous curvature of flat lattices

Our data support the constant area model for clathrin curvature in the transition between domes and spheres across the eight cell lines tested. Yet, we cannot assign this model to the preceding stage of flat to domed transition. Thus, we directly tested if flat structures could curve at the plasma membrane. Past data has indicated that clathrin in unroofed cells can spontaneously curve at neutral pH.^34^ We further tested this phenomenon quantitatively and across many conditions (Fig. 3). HeLa cells were unroofed but fixed after resting in neutral HEPES buffer for increasing amount of time. Figure 3 shows that under these conditions at the 20-minute time point a significant decrease in the density of flat clathrin is evident with a corresponding increase in spherical clathrin. We tracked these changes at 0, 2, 5, 10, 20 minutes post-unroofing (Fig. 3 e-g). During that time, the area occupied by flat clathrin decreased 8-fold from 2.1 ± 1.8 % to 0.26 ± 0.16 % while the area occupied by spherical structures increases 7-fold from 0.29 ± 0.18% to 2.1 ± 11.2%. At 20 minutes, the flat structures that did remain were small (55.6 ± 12 nm at 20 min vs. 100.6 ± 35.1 nm at 0min; 3.3-fold area difference). Perhaps these lacked enough triskelia to curve. The remaining domes were also small (effective radius 64.2 ± 12.2 nm at 20 min vs. 87.2 ± 17.7 nm at 0 min). However, the spheres formed after unroofing were roughly the same size as naturally created spheres (effective radius 75.0 ± 15.8 nm at 20 min vs. 69.4 ± 11.4 nm at 0 min). These data are consistent with lattices relaxing to an equilibrium curvature. Importantly, the total amount of clathrin did not decrease during the treatment (Fig. 3g). This is supported by a fluorescence microscopy where live HeLa cells expressing fluorescence clathrin light chain were imaged before and after unroofing. Clathrin structures remained on the surface and lost no more than 30% of their total fluorescence even after 20 minutes (Fig. S3). Together, these experiments indicate that flat clathrin can curve in native membranes. Importantly, clathrin can curve spontaneously without the recruitment of curvature-inducing factors (proteins) or added energy (ATP or GTP). In opposition to previous models, these data show that curvature on the membrane can occur without bulk cytoplasmic clathrin exchange.

**Figure 3.**
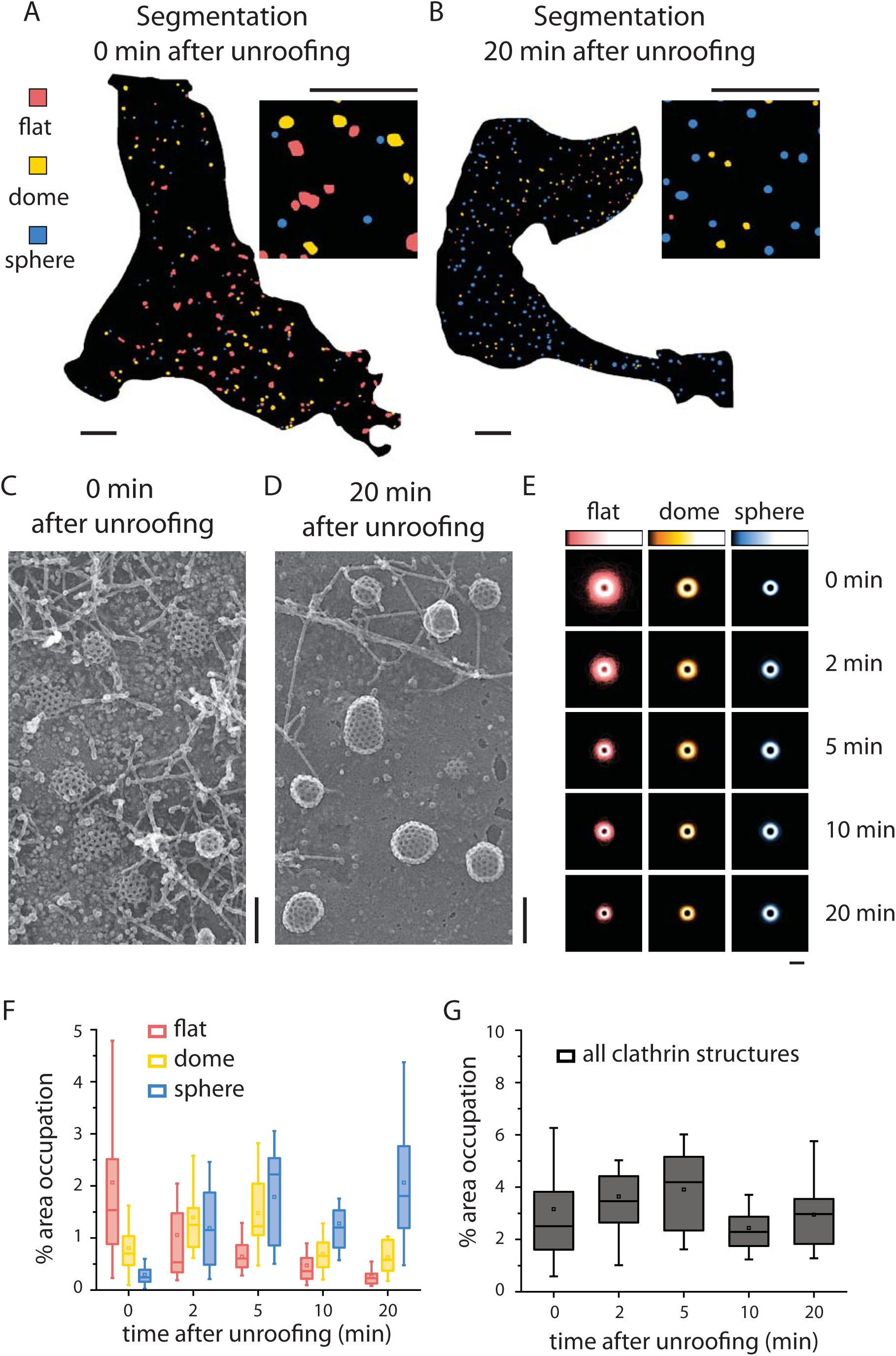
Flat clathrin structures spontaneously curve with no added factors or energy. (A) Segmentation of a HeLa cell fixed during unroofing as in Figs 1-5. (B) Segmentation of a HeLa cell unroofed, placed in fresh buffer, and fixed 20 minutes later. Insets are shown at higher magnification. (C) An example of a platinum replica 0 minutes after unroofing and (D) 20 minutes after unroofing. (E) Binary outlines of each structure were centered and summed for each segmentation category as in Fig. 1i. The relative intensity scale for each category is shown. (F) The percent of membrane area occupied by flat, dome, and sphere structures is shown for 0, 2, 5, 10, and 20 minutes after unroofing. (G) The sum of flat, dome, and sphere membrane area occupation shows total clathrin area occupation. 0 min, N_flat_=2585, N_dome_=1438, N_sphere_=936, N_cells_ = 38; 2 min, N_flat_=852, N_dome_=1216, N_sphere_=1066, N_cells_ = 14; 5 min N_flat_=608, N_dome_=1194, N_sphere_=1183, N_cells_ = 9; 10 min, N_flat_=705, N_dome_=859, N_sphere_=1059, N_cells_ = 14; 20 min, N_flat_=456, N_dome_=842, N_sphere_=1885, N_cells_ = 14. For box plots, box is interquartile range, center line is median, center circle is mean, whiskers include all data excluding outliers. 0 min timepoint includes HeLa data from Fig. 1. Scale bars for A-B are 2 µm. Scale bars for C-E are 200 nm.

### Flattening Forces

If clathrin can spontaneously bend, what keeps it flat in living cells? To answer this question, we tested if curvature of clathrin could be modulated in living or unroofed cells. Because clathrin plaques are less prevalent on unadhered surfaces,^28, 35^ we reasoned that physical adhesion to the substrate could impart a flatting force. β5-integrins have been shown to localize to clathrin plaques.^19, 24^ Figure 4 shows that disrupting the connection between β5-integrins and the extracellular matrix with the drug cilengitide (CTA) caused a 2-fold loss of flat structures in HeLa cells (0.96 ± 0.35% with CTA vs. 2.1 ± 1.8% membrane occupation in control). This coincided with a 2-fold increase in domed structures and a 3-fold increase in spherical structures (1.7 ± 0.51% domes, 0.76 ± 0.38% spheres in CTA vs. 0.80 ± 0.53% domes, 0.29 ± 0.18% spheres membrane occupation in control). There was, however, not a complete loss of flat structures after the drug treatment. Either the drug did not completely disrupt β5-integrin or not all flat structures are kept flat with adhesion through integrins. While this drug had a major effect on clathrin structure, it is possible that additional forces could contribute to the creation of flat structures.

**Figure 4.**
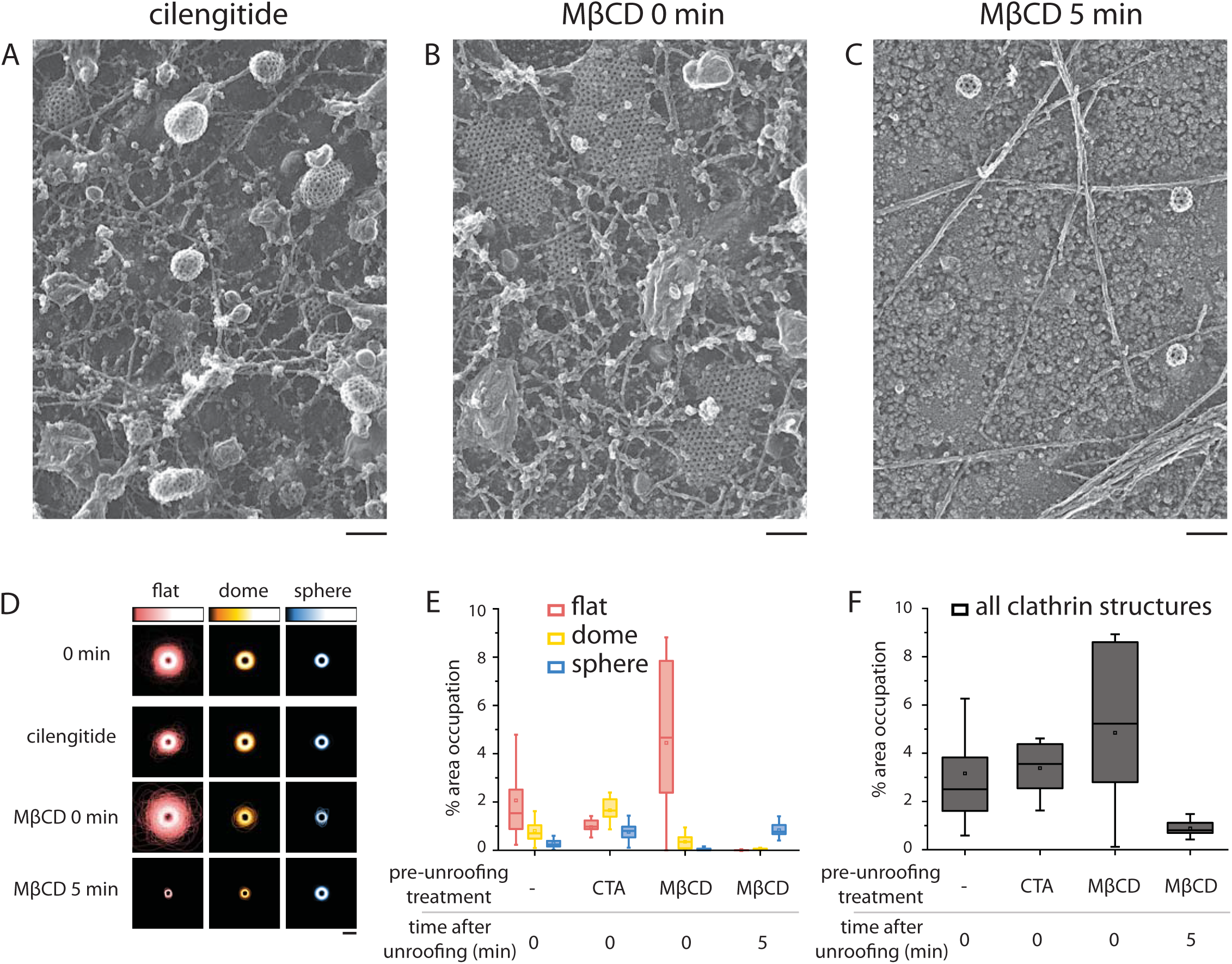
Membrane adhesion and cholesterol content can modulate the amount of flat clathrin but disassembly of flat clathrin dominates over curvature on a cholesterol depleted membrane. HeLa cells were treated with (A) cilengitide (CTA) or (B) methyl β cyclo dextrin (MβCD) for one hour prior to unroofing/fixation. (C) Cells treated with MβCD were alternatively unroofed and fixed 5 minutes later. (D) Binary outlines of each structure were centered and summed for each segmentation category as in Figure 1i. (E) The percentage of membrane area occupation from flat, dome, and sphere structures is shown as a box plot. The 0 min timepoint data is shown for reference from Figure 3. (F) The sum percentage of membrane area occupation is shown as a box plot. For E-F, box = interquartile range, center line = median, center circle = mean, whiskers include all data excluding outliers. Scale bars for A-C are 200 nm. No treatment, N_flat_=2585, N_dome_=1438, N_sphere_=936, N_cells_ = 38; cilengitide, N_flat_=729, N_dome_=1218, N_sphere_=673, N_cells_ = 13; MβCD, 0 min, N_flat_=597, N_dome_=141, N_sphere_=28, N_cells_ = 11; MβCD, 5 min, N_flat_=18, N_dome_=68, N_sphere_=766, N_cells_ = 9. Untreated HeLa data is from Figure 3 and shown for reference.

Therefore, we tested the hypothesis that flat structures could be controlled by other forces such as membrane stiffness. To test this, HeLa cells were cholesterol-depleted with methyl-β-cyclodextrin (MβCD). Cholesterol depletion is thought to stiffen the membrane, making membrane fluidity and thus vesicle formation more difficult.^36^ As previously suggested,^37, 38^ we observe an increase in flat clathrin structures (2-fold, 4.5 ± 3.2% with MβCD vs. 2.1 ± 1.8% membrane occupation in control) and a corresponding decrease in domed (2-fold, 0.35 ± 0.33% with MβCD vs. 0.80 ± 0.53% membrane occupation in control) and spherical structures (7-fold, 0.04 ± 0.05% with MβCD vs. 0.29 ± 0.18% membrane occupation in control) (Fig. 4b,d-f; 0 min refers to fixation during unroofing). This suggests modification in membrane stiffness can prevent a flat to curve transition. Next we tested if spontaneous curvature would not occur in cells treated with MβCD. Figure 4e shows that five minutes after unroofing, MβCD-treated cells still exhibited an increase in spherical structures (0.83 ± 0.31% with MβCD 5min vs. 0.04 ± 0.05% membrane occupation in MβCD 0 min, 5 min refers to fixation five minutes after unroofing). Yet, the total amount of clathrin on the membrane after unroofing dropped over 5-fold (0.88 ± 0.35% with MβCD 5min vs. 4.8 ± 3.2% membrane occupation in MβCD 0 min) (Fig. 4f). This could indicate that strained flat lattices are not tightly networked together. In the case that they cannot ease this strain by bending, the triskelia might then disassemble.

### Flat lattices are preloaded with pentagons

To determine if flat lattices are loosely networked, we analyzed the detailed images of lattices from platinum replica EM. As previously reported,^33^ flat lattices frequently have curved pits budding from them. Anywhere from 9.5% (L6) to 39% (MCF7) of the observed flat structures had a curved structure immediately adjacent (Fig. 5a, Fig. S2g-h). This is surprising because it indicates that the forces that keep structures flat must be very localized. Further, it showcases that lattices can cleave in a coordinated way into subzones of activity. Figure 5b-c shows that in striking contrast to common models of flat lattices, clathrin regions commonly contained breaks, imperfections, and extended fissures (Fig. 5b-c). Modeling in Figure 5d shows that this would be impossible if triskelia were centered at each vertex. Specifically, four lattice vertices must be missing a triskelion to accommodate one broken segment or strut. Therefore, we support a model where flat lattices are not saturated with triskelia.^39^ Beyond these breaks, holes in the lattice differed in size and were not organized according to regular hexagonal lattice packing. Such packing would maintain equal sides and angles and linear arrays of holes. These observations support a much more disordered conformation of flat clathrin lattices than previously proposed. We propose that this disorder could support the transitions needed for lattice curvature.

**Figure 5.**
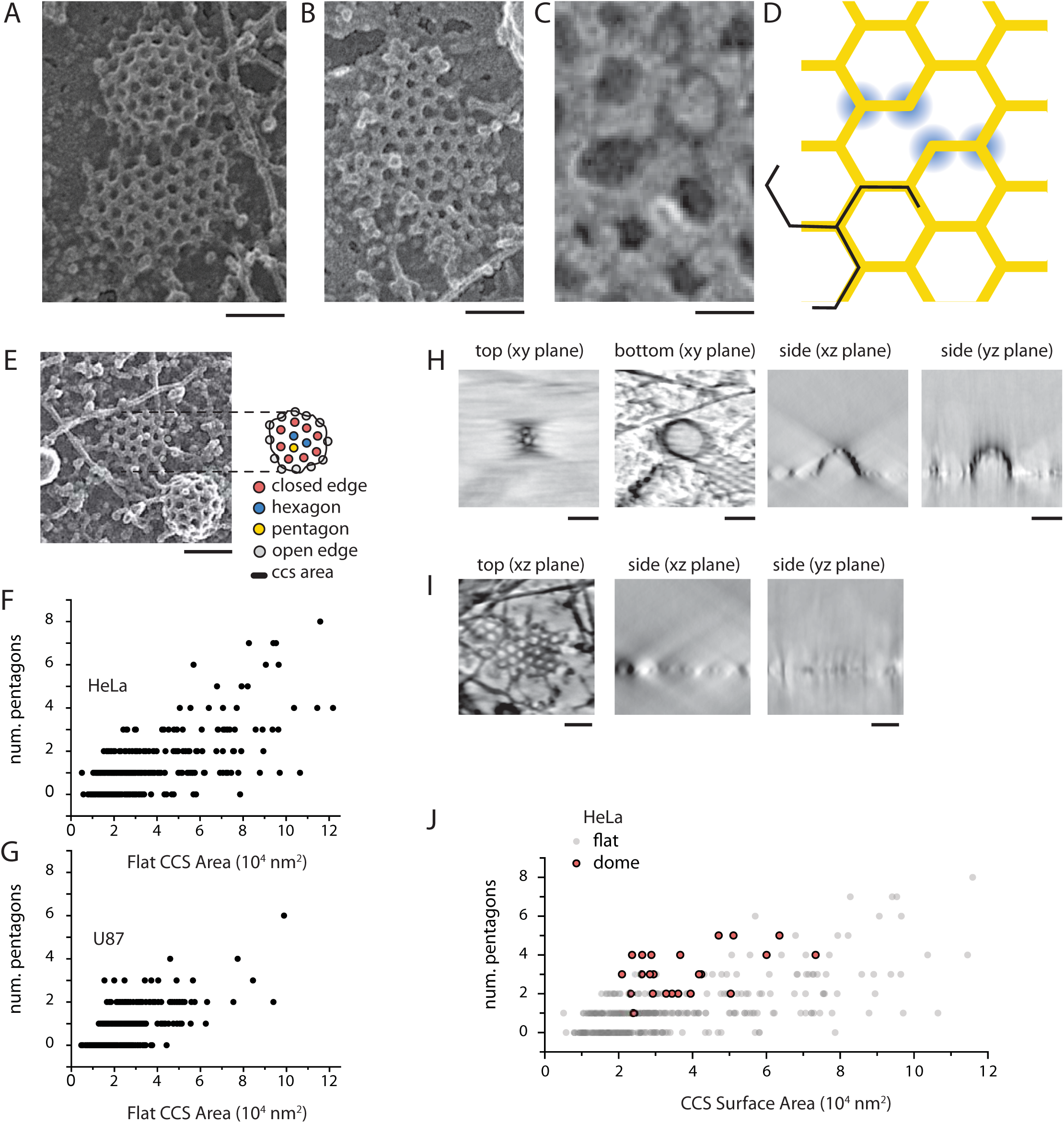
Pentagons are frequently found in flat structures. (A) A representative image of a dome budding immediately adjacent to a flat structure. (B) A representative image of flat clathrin with a break in the center of the lattice with a zoomed in view shown in (C). (D) A schematic showing a lattice with a single break (yellow). For the break to exist, the lattice cannot be fully occupied. Triskelia like the one shown in black must be missing from the highlighted vertices (blue), suggesting flat structures are not fully saturated. (E) Polygons in flat structures were labelled as open edge, closed edge, hexagons, and pentagons as shown. (F) The number of pentagons in flat structures is plotted vs the lattice area for HeLa cells and (G) U87 cells. (H) A tomogram of a domed clathrin structure from a platinum replica is shown in four different planes. (I) A tomogram of a flat clathrin structure from a platinum replica is also shown. (J) The number of pentagons counted in domed structures from tomograms of HeLa cells is plotted vs surface area. The flat data from B are shown again for reference. Scale bars from A-B, E, H, I are 100 nm. The scale bar C is 25 nm.

Pentagons are necessary to form an enclosed vesicle in a regular hexagonal lattice. Disorder could allow pentagons to exist prior to curvature and lessen the need for gross lattice rearrangements. Therefore, we tested if flat lattices contain pentagons. We counted the number of pentagons in flat lattices of HeLa cells (Fig. 5e-g). Pentagonal holes in clathrin lattices were confirmed by counting the number of surrounding holes. Thus, holes at the edge of the structure were not included in this analysis (Fig. 5e). Figure 5f shows that pentagons exist in most flat structures, even very small ones (Fig. 5f). As flat structures got bigger, an increased number of pentagons were present. The same trend was observed in U-87 cells (Fig. 5g). To see how this pentagon density compared with the density in domed structures, we used PREM tomograms to count the pentagons in domes (Fig. 5h-j). While pentagons were on average slightly more common in domed structures, many flat structures had the same surface density of pentagons (Fig. 5j). Pentagons in a regular flat hexagonal lattice is geometrically impossible. Therefore, it is important to note here, that hexagons and pentagons in these structures are not regular with equal length sides and angles. Indeed, the polygons in these lattices frequently appear skewed with struts of varying lengths. Additionally, our analysis classified holes into an “other” category that were neither pentagons nor hexagons and were difficult to identify. Very large irregular holes with more than six sides fell into this category, their identity often obscured by increased breakage at these spots. Thus, the clathrin lattice in the flat state is flexible. In a flexible lattice, spontaneous curvature can occur without the addition of triskelia or large scale reorganization. The thickness of platinum, however, in these and previous data limited our ability to directly observe molecular-scale lattice organization.

### Flat lattices are flexibly assembled with loosely connected triskelia

To overcome this limitation, we used cryoET to observe the molecular-scale interactions between single triskelia in flat lattices on cell membranes. Flat, domed, and spherical structures were observed but precise segmentation of curved lattices was not possible due to the missing wedge feature of cryoET and the fixed geometry of our samples. Flat structures were segmented by manually determining the positions of vertices in three-dimensions. Segmentation-guided isosurfaces aided in 3D visualization and confirmed flatness in these structures (Fig.6a-c, Fig. S4, stack shown in Supp. Movie 1). Strut-to-strut angles and strut lengths were extracted from the segmentation (Fig. 6d-e). The average angle within a flat lattice made of triskelia is geometrically defined to be 120 degrees which is what we calculate when we constrain the angles to two dimensions (120.0 +/− 13.3 degs). When the angles are measured in three-dimensions, the average angle is expected to decrease with bending. However, we observe no substantial decrease when the angles are measured in three-dimensions (119.3 +/− 13.0 degs, mean +/− SD) consistent with these being flat lattices. The average strut length measured at 18.4 +/− 2.0 nm is also consistent with previous reports in clathrin baskets.^40, 41^ However, a high degree of flexibility is indicated in these data as the angles ranged from 75 to 157 degrees and strut lengths range from 8.2 to 24.1 nm. These data again highlight the flexibility of the subunits in these structures.

**Figure 6.**
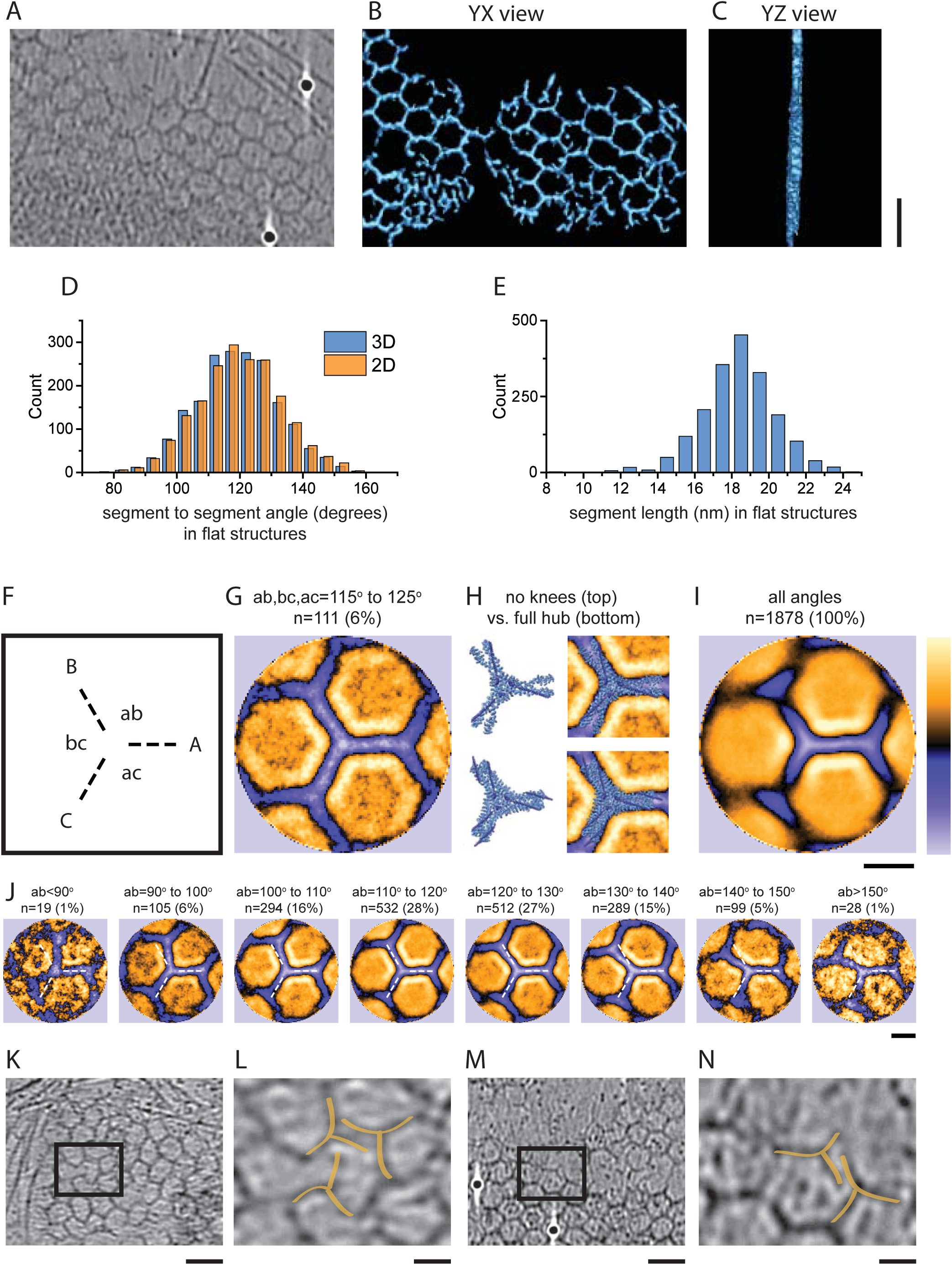
Cryo-electron tomography of flat clathrin structures show irregularity and loosely connected triskelia. (A) An XY-slice of an example tomogram of a flat clathrin structure in HeLa cells, and its segmentation guided isosurface (B, XY view; C, XZ view). (D) A histogram of the 3D and 2D (XY slice plane) angles between struts in flat structures. (E) A histogram of the length of struts between vertices in the flat structures. (F) Vertices with three struts radiating outward were used for averaging. Each vertex of interest was centered, and rotated to extend the strut of interest along A as shown. (G) When all angles are filtered between 115° and 125°, to obtain only the most symmetric vertices, 6% of the data remains. The average is shown. (H) The clathrin basket consensus hub (PDB:6SCT)^14^ is shown with the 1^st^ degree neighbor triskelia removed (no knees), or the full structure. The structures are shown with the average from G to illustrate the missing knee density in the average. (I) The average from all struts within a 3-way vertex is shown. (J) Each segmented strut was placed into cohort based on the angle counterclockwise (ab). The averages from each cohort are shown. A white dashed line is shown to reference 120° regular angles as in F. (K, M) Two examples of tomograms are shown and one of several examples of broken struts is highlighted and (L, N) magnified. The orange highlights clearly separated triskelia legs within lattice struts. The scale bars for A-C, K and M are 50 nm. The scale bars for F-J, L and N are 15 nm. For F-J, the lookup table for color scaling is shown to the right of I.

To obtain an average vertex structure, the vertices were centered with translation and the tomograms were rotated to align the strut of interest (Fig. 6f-j; strut A). Data were filtered to only include struts within highly symmetric vertices with angles within 115 and 125 degrees leaving only 6% of the original data (111 of 1878). The average of these select data is mostly consistent with the single particle cryoEM determined consensus hub of clathrin triskelia assembled into baskets.^14^ However, our average exhibited a narrower density immediately surrounding the vertex where the previously determined basket consensus hub predicts the location of the knees from the neighboring triskelia (Fig. 6g-h). Possibly, flat lattices have a low occupation of direct neighbors. This would be expected from the broken struts seen in our platinum replica data (Fig. 5 b-d). Alternatively, it could suggest that the knee is heterogeneous or placed in a different confirmation in flat structures.

When all the tri-connected struts are included in the tomogram average, the variability in angles prevents a clear structure aside from the aligned strut (Fig. 6i). Therefore, we separated the data into angular cohorts based on the angle counter clock-wise to the strut of interest (Fig. 6f,i; angle ab). Here, we can see that flat structures contain much more angular variability than would be expected from the single particle cryoEM basket consensus hub.^14^ In clathrin baskets, the regular interlacing and bonding between subunits imparts a highly ordered structure. But the inherent flexibility of flat lattices limits our ability to make atomic-scale generalizations about lattice structure. Therefore, we measured objects in single tomograms to determine the molecular traits that accommodate assembly. At several locations, we can clearly detect separated or broken struts (Fig. 6k-l, Supp. Movies 2-3). Legs of neighboring triskelia can be spaced by distances of up to 5 nm. Based on these distances, we suggest that the shape of clathrin triskelia and the positioning of inter-triskelia binding sites do not permit full networking within an extended flat lattice. Angular variability and disconnected struts could explain how pentagons can be present and common in these lattices. These data suggest that spontaneous curvature might occur by strengthening inter-triskelia connections rather than gross lattice reorganization, addition or exchange of clathrin subunits, and energy dependent lattice remodeling. A full model incorporating our findings is presented in Figure 7.

**Figure 7.**
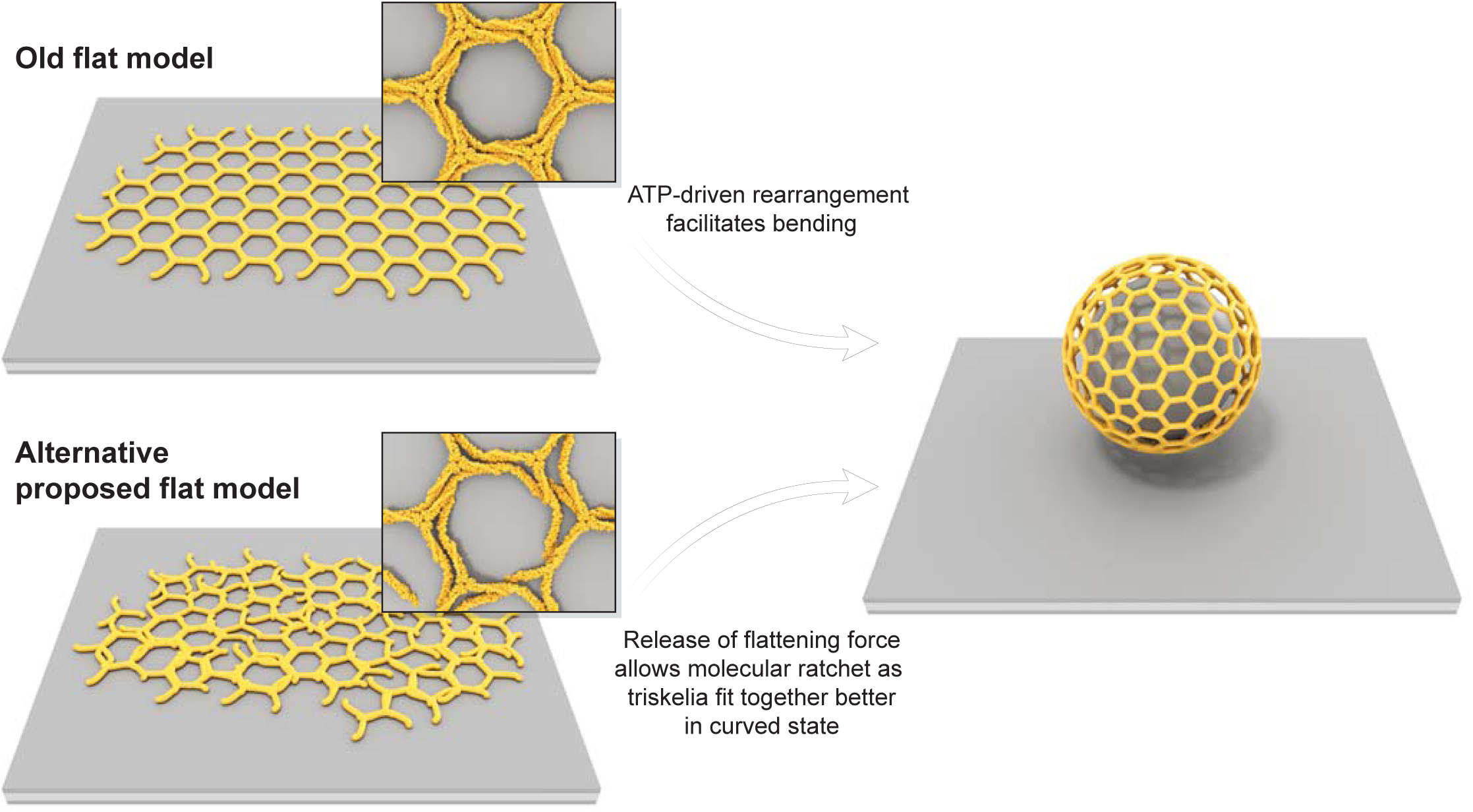
A revised model for the curvature of flat clathrin lattices. In the current model, flat clathrin is depicted as a flat version of cryoEM clathrin basket structures. The triskelia are tightly connected, interlaced, and fully saturated. Therefore, energy is required to exchange triskelia with cytoplasmic triskelia and remodel clathrin into a curved structure. In the new model, triskelia in flat lattices are not fully networked to one another and lattices are not fully saturated. The natural flexibility and loose networking allows pentagons to be already present in this flat state. A flattening force prevents triskelia from assembling in their preferred curved state. Upon removal of the flattening force, triskelia spontaneously move into a more preferred state as in a Brownian ratchet.

## Discussion

We have explored the spatial and structural patterns of clathrin mediated endocytosis in mammalian cells at the nanoscale. We show that in eight commonly used cell lines, lattices maintain a near constant surface area in the latter half of the curvature process. Further, flat clathrin lattices can spontaneously bend without the addition of triskelia, protein factors, enzymes, or energy. The amount of flat clathrin in living cells can be modulated by disrupting integrin-based adhesion or stiffening the membrane with cholesterol removal. Stiff membranes caused increased clathrin loss during spontaneous curvature suggesting that these structures are under strain. The structure of flat clathrin is suited for spontaneous curvature because it is preloaded with pentagons, not fully saturated with subunits, and is not fully constrained into a rigid highly-interwoven lattice. Collectively, these data support major changes to the current model of energy-dependent clathrin exchange and lattice remodeling during vesicle formation at the plasma membrane.

Our data support a model where tight lattice assembly is prevented during the initial flat growth phase of endocytosis. Removal of a multi-factored flattening force allows for inter-triskelia connections that favor the curved state (Fig. 7). Here, the driver for clathrin curvature is contained in the inherent properties of the lattice and its associated proteins.

To date, flat clathrin has generally been modeled as a planar, tightly interwoven, strictly hexagonal version of the consensus cryoEM clathrin basket structure.^13, 14^ This presents a substantial energetic barrier to lattice bending. We confirm that flat structures are common. Previous work with electron, live cell fluorescence, and correlative microscopy has shown that in some cell lines (SK-MEL-2, BS-C-1)^9, 12^ flat clathrin curves. Further, in these cells curvature can occur at the onset, during, or after complete recruitment of clathrin to the membrane.^8^ Here, we show that this mode of endocytosis is not unique to these two cells lines. Specifically, all eight cell lines studied show near full lattice recruitment by the hemispherical state. We also show that, contradictory to most interpretations of the constant area model of endocytosis, it is possible for these flat lattices to bend into spherical buds without the addition of more triskelia or energy.

One argument for the constant area model has been that triskelia are always exchanging with cytoplasmic clathrin. This exchange is thought to enable the rearrangements that would be needed to introduce pentagons into hexagonal flat lattices.^12, 42^ The ATPase HSC70 and its co-chaperone auxilin, which remove the clathrin lattice after scission, have been implicated in exchange during pit maturation.^42, 43^ In support of this idea, clathrin structures in live cells exhibit a 25-86% exchange of clathrin with 2-25 s half times of recovery (t_1/2_) when measured with fluorescence recovery after photobleaching (FRAP).^12, 42, 44^ Clathrin exchange is inhibited by ATP-depletion or auxilin mutations. But critics point out that auxilin is not present during all stages of coat assembly preceding and up to the moment of scission. Mutating or knocking down these proteins in an effort to inhibit clathrin exchange could instead cause a build-up of cytoplasmic coated vesicles that are easily misinterpreted as membrane bound endocytic structures.^45^ Further, the time scale and total amount of recovery in FRAP experiments can be consistent with clathrin growth rather than exchange. Lastly, no clathrin FRAP experiment exhibits recovery over 100%. This over-recovery would be expected in a growing structure experiencing complete dynamic exchange. Surprisingly, our data instead show that flat clathrin can bend to form vesicles in the absence of ATP or clathrin exchange. During spontaneous curvature, clathrin structures lost no more than 30% of clathrin triskelia. Part of this loss could be due to photobleaching and thus the effect is clearly an overestimate of clathrin loss. ATP-dependent clathrin exchange is likely critical for vesicle uncoating but might not occur during the assembly and curvature of the lattice. Our data support this model.

A previous reconstitution experiment demonstrated that flat clathrin can polymerize on a membrane at 4°C and then spontaneously bud at 37°C.^46^ Thus, at physiological temperatures, curved clathrin is favored. Even in the case of increased membrane tension, reconstitution experiments at physiological temperatures create shallow domes rather than extended flat lattices.^47^ Furthermore, many clathrin accessory factors sense or facilitate membrane curvature thus making curvature even more favored.^48^ As an example, epsin is known to create membrane curvature by two different mechanisms; amphipathic helix insertion,^1, 49^ and steric asymmetry induced by an unstructured bulky cytosolic domain.^50, 51^ Yet, in live cells, clathrin can grow flat (Figs. 1-2). And flat structures in HeLa cells are loaded with an abundance of epsin and other membrane bending factors.^52^ Therefore, our model proposes that flat structures require a significant flattening force to overcome both clathrin’s preference for curvature and these curvature-inducing or stabilizing factors. We suggest that upon release of this flattening force, pre-loaded curvature inducing proteins help clathrin relax into its preferred state of curvature.

There does not seem to be one flattening force that causes the formation of flat structures. Here, we manipulate the flattening force in two different ways: 1) disrupting integrin connections to the extracellular matrix, and 2) removing cholesterol. Cilengitide selectively inhibits αVβ5-integrin and αVβ3-integrin connections with the extracellular matrix.^53^ αVβ5-integrin has been shown to localize to flat clathrin structures.^19, 24^ We show that removal of this connection vastly decreases the amount of flat clathrin in HeLa cells. In fact, the relative representation of flat structures in HeLa cells after cilengitide treatment looks similar to the relative amount of flat clathrin in cell lines not generally considered to have plaques (BS-C-1, PC-12) (Figs 2,4). Integrin clathrin adhesion sites could account for the difference in morphology and amount of flat clathrin between these different cell types. Recently, alternative splicing of clathrin heavy chain in myotubes has been shown to decrease plaques to a similar degree indicating a possible interaction site controlling integrin-based clathrin adhesions.^54^ Due to technical limitations of our imaging modalities we are limited to studying adherent membrane surfaces. It is possible that the density of flat clathrin on the apical side of cells^35^ could show less diversity among cell lines. Future work to study non-adherent surfaces, non-adherent cells, or adherent surfaces in 3D tissues systems at this resolution and breadth will require the development of substantial new technologies. Aside from these limitations, we feel that these basic aspects of clathrin assembly are universal.

Beyond clathrin adhesion and membrane stiffness, other flattening forces are likely present at the plasma membrane. In non-adherent membranes, the actin cortex could block lattice curvature. Indeed, some flat structures in cryoEM exhibited long clathrin fibers extending over top of the lattice (Fig. S4). During unroofing, some membrane tension is released, the actin cortex is removed, and the bond between integrins and adaptors could be severed. In a living cell, however, it is unclear how the removal of these factors could be locally coordinated at a single structure. In some cases, actin likely plays an important local role in lessening membrane tension.^55-58^ But it is curious that curved clathrin structures frequently exist immediately adjacent to flat structures (Fig. S2g). If tension is released it must be a very local event. Future work will look for signaling events that might allow local changes to curvature at the scale of a single organelle.

Other local events could regulate flat clathrin assembly. These could include the loss or recruitment of a protein factor, conformational changes, or signaling event (phosphorylation) within flat structures that prevent tight binding of triskelia. Interestingly, clathrin light chain can be phosphorylated or manipulated by cargo and adaptor proteins.^59-61^ Alternatively, tight lattices likely assemble with residue-specific interactions that are geometrically disallowed in flat structures^16^. This is supported by cryoEM of basket structures where even at slightly lower curvature, geometric constraint results in fewer inter-triskelia bonds.^14^ We suggest that triskelia in flat structures have fewer binding sites, creating lower avidity and a loosely connected lattice.

In summary, we present a new universal model where highly flexible triskelia assemble into a flat lattice which lacks tight inter-triskelion bonds. This type of lattice assembly allows flat structures to be preloaded with pentagons minimizing the need for gross rearrangement to the lattice during curvature. In our cryoET data, this loose assembly is visible as variations in strut length, strut-to-strut angles, leg separations and broken lattices. This is consistent with visibly broken lattices observed in platinum replica data. In agreement with a recently introduced model,^39^ our ultrastructural analysis suggests flat clathrin are also not fully saturated with triskelia at each vertex. With the loss of a flattening force, and the aid of curvature inducing proteins, the lattice spontaneously assumes the curved state to form a vesicle. Future work will be needed to determine how the dozens of local protein factors, cargo, and signaling events are coordinated to regulate and control this dynamic complex assembly at the plasma membrane.

## Material and Methods

### Cell Culture

PC12-GR5 (rat adrenal gland, pheochromocytoma) cell line originated from Rae Nishi’s lab (OHSU). They are a common cell line for the study of exocytosis. All other cell lines were all purchased through ATCC. 3T3-L1 (ATCC^®^ CL-173^™^, mouse embryo fibroblast), BS-C-1 (ATCC^®^ CCL-26^™^, *Cercopithecus aethiops*, kidney epithelial), HeLa (ATCC^®^ CCL-2^™^, human cervix epithelial, adenocarcinoma), L6 (ATCC^®^ CRL-1458^™^, rat skeletal muscle, myoblast), MCF7 (ATCC^®^ HTB-22^™^, human mammary gland, epithelial, adenocarcinoma), RAW264.7 (ATCC^®^ TIB-71^™^, mouse, Abelson murine leukemia virus-induced tumor, monocyte/macrophage), U-87 MG (ATCC^®^ HTB-14^™^, human, brain, epithelial, likely glioblastoma). 3T3-L1, BS-C-1, HeLa, L6, U-87 cell lines were grown in DMEM (no phenol red, Thermo-Fisher, Gibco™, 31053036) with 10% fetal bovine serum (Atlanta Biologicals, S10350), 1:100 dilution of 100x Glutamax (Thermo-Fisher, 35050061), 1:100 dilution of 100 mM sodium pyruvate (Thermo-Fisher, Gibco™, 11360070), with or without 1:100 dilution of Penicillin-Streptomycin (Thermo-Fisher, Gibco™, 5,000 U/mL). MCF7 and RAW264.7 cell lines were grown in DMEM (high glucose, pyruvate included, Thermo-Fisher, Gibco™, 11995073), 10% fetal bovine serum (Atlanta Biologicals S10350), and 1:100 dilution of Penicillin-Streptomycin (Thermo-Fisher, Gibco™, 5,000 U/mL). PC-12 cells were grown in DMEM (high glucose, pyruvate included, Thermo-Fisher, Gibco™, 11995073) with 5% fetal bovine serum (Atlanta Biologicals, S10350), 5% horse serum (Atlanta Biologicals, S12150), and 1:100 dilution of Penicillin-Streptomycin (Thermo-Fisher, Gibco™, 5,000 U/mL). RAW264.7 cells were grown on Fibronectin coated coverslips (Neuvitro, GG-25-1.5-fibronectin). All other cell lines were grown on lab-coated (Sigma, P4832-50ML) or commercially coated Poly-Lysine (Neuvitro, GG-25-1.5-PDL) coverslips.

### Platinum replica electron microscopy

Cells were rinsed briefly in unroofing buffer (30mM HEPES, 70mM KCl, 5mM MgCl2, 3mM EGTA at pH 7.4) immediately prior to unroofing. Cells were unroofed by physical disruption of the plasma membrane either by syringe or sonication. With sonication, the coverslip was moved to unroofing buffer containing 0.5% paraformaldehyde (Electron Microscopy Sciences, 30525-89-4) immediately prior (within 10 seconds) to two 400 ms sonication pulses from a Branson Sonifier 450 (VWR International, 47727-492) with a 1/8” tapered microtip (VWR International, 33996-163). The coverslips were then moved immediately into unroofing buffer containing 2% glutaraldehyde (Electron Microscopy Sciences, 16019). With syringe, coverslips were sprayed briefly with unroofing buffer containing 2% paraformaldehyde using a 22-gauge needle held vertically within 2 inches of the coverslip. The syringe was moved around the coverslip during the spray. The coverslip was immediately moved to fresh unroofing buffer containing 2% glutaraldehyde. Cells were kept in glutaraldehyde at 4°C until ready for staining and dehydration; less than two days. Staining, dehydration, drying and platinum coating were performed as described previously.^31, 52^ Briefly, the cells were stained with 0.1% tannic acid and 0.1% uranyl acetate and dehydrated slowly through a series of increasing ethanol concentration to 100% ethanol. The coverslips were then dried with critical point drying and rotary shadowed with platinum and carbon. The replicas were released from the coverslip with hydrofluoric acid and placed onto a TEM grid for imaging. TEM imaging of replicas was performed on a JEOL 1400 equipped with SerialEM freeware^62^ for montaging and tomograms.

### Spontaneous curvature and drug treatments

Spontaneous curvature experiments were performed similarly to normal unroofing experiments. However, after unroofing, the coverslips were moved to fresh unroofing buffer for the specified time prior to fixation. Cilengitide (10 µM, Sigma-Aldrich, SML1594) or methyl-β-cyclodextrin (9.5 µM, Sigma-Aldrich C4555) were added to growth media at normal growth conditions for one hour prior to unroofing and fixation.

### Analysis of platinum replicas

Montages were stitched together using IMOD.^63^ Each montage was manually segmented in imageJ^64^ by outlining the edge of the membrane, flat clathrin structures (no visible curvature), domed clathrin structures (curved but can still see the edge of the lattice), and sphere clathrin structures (curved beyond a hemisphere such that the edge of the lattice is no longer visible). Membrane area fraction was defined as the sum of area from clathrin of the specified subtype divided by the total area of the membrane. Effective radius was defined as 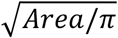 and was used as a simple way to depict area on a linear scale as perceived by the eye. Percent area representation was the sum of area from clathrin of the specified subtype divided by the sum area of all the clathrin on the membrane. “Area” refers to two-dimensional image area. “Surface Area” refers to the area of the 3D clathrin lattice. For pentagon counting (Fig. 5), a custom Matlab program was used to automatically detect holes in the clathrin lattices. These holes were manually designated as edge, pentagon, hexagon, and other. Manual removal or addition of holes was performed as needed. Surface area for flat structures was determined by outlining segmentation (as for Figs. 1-2). Surface area for domes was calculated based on height and width of the structure assuming a hemisphere.

### Fluorescence imaging of unroofing (supplemental)

HeLa cells were transfected and plated onto Poly-Lysine coated coverslips (Neuvitro, GG-25-1.5-PDL) one day prior to imaging. Transfection was performed following the Amaxa Cell Line Nucleofector Kit R protocol. Cells were transfected with mCherry – clathrin light chain B (Addgene Plasmid #55019) and EGFP – clathrin light chain A^52^ plasmids. Cells were moved into unroofing buffer immediately prior to the imaging series. Acquisitions were 50 time points with a time point every 30 seconds (0.033 Hz).

During that time, up to eight regions of interest were imaged with four frames: 488 nm total internal reflection in ring formation (ring TIRF)^65^, 561 nm ring TIRF, 488 nm epifluorescence, and 561 nm epifluorescence. Each frame was 100 ms exposure. Unroofing was performed by syringe between the sixth and seventh time point of the acquisition. Cells were imaged on an Olympus IX81 inverted microscope equipped with a 3I Vector TIRF launch with two fiber optic inputs and galvo steering. Images were projected onto an Andor Ixon EMCCD through a DualView (BioVision Technologies) image splitter. 100 nm tetraspeck beads were imaged before each session and used to align the red to the green images using a projective transform. Images were analyzed in ImageJ^64^ using the Time Series Analyzer plugin, selecting 20 clathrin puncta of interest per cell and a background region on interest nearby. The intensity of each region over time was background subtracted. Intensity over time of each clathrin structure was normalized to the frame after unroofing. We present the average and standard deviation of the background subtracted and normalized TIRF signal over time across all clathrin puncta. Epifluorescence was only used to assess effective unroofing.

### Preparation of cryoEM samples

Quantifoil gold 300-mesh grids with R2/1 carbon hole size (SPI) were sterilized with a UV light for 40 minutes. Grids were then coated for 1 hour with a 1:40 dilution of fibronectin (Sigma) in water. Cells were densely plated on grids and allowed to settle on the grids for one hour before removing excess cells and placing the grid in fresh growth media. After one day of growth, grids were prepared for cryoEM. Freshly removed from the incubator, each grid was placed onto plunging tweezers, rinsed briefly with unroofing buffer (above), placed against the lid of a 6-well dish, and unroofed by spraying with 0.5% paraformaldehyde in unroofing buffer. The edge of the grid was blotted. 2 µL of unroofing buffer and 2 µL of 10 nm BSA gold tracer solution (Electron Microscopy Sciences) was added to the grid prior to placing in a Leica plunge freezer. In the plunge freezer, the grid was blotted from the back for 3 seconds immediately prior to plunging into liquid ethane. The cells were plunged within 20 seconds of unroofing.

### Acquisition of cryoEM data

Vitrified grids were imaged on an FEI Titan Krios electron microscope operating at 300 kV at 26000x projecting onto a Gatan GIF Quantum K2 camera in electron counting mode. The resulting pixel size was 5.37 Angstroms. Tilt series were acquired from 0 degrees to −50 degrees and then 0 to 50 degrees in 2 degree increments. Six seconds of acquisition was performed at each angle over 12 frames. Total electron exposure throughout the tilt series was below 130 electrons/Å^2^. Defocus was between −6 µm and −10 µm. Tilt series were acquired with SerialEM.^62^

Motion correction was performed with the alignframes function in IMOD.^63^ Tomograms were further processed using etomo in IMOD. The tilt series were pixel defect corrected and aligned with gold particle tracking. Tomograms were reconstructed using the simultaneous iterative reconstruction technique (SIRT) with six iterations. The single tomogram slices shown in Fig. 6k-n were filtered using the imageJ plugin DenoisEM^66^ Tikhonov deconvolution (*λ* = 15, *σ* = 3, and *N* = 30 iterations).

### Analysis of cryoEM data

18 flat structures were identified by eye across ten tomograms. Each structure was manually segmenting in 3DMOD (IMOD) by identifying the 3D coordinates of vertices. A custom Matlab (The Mathworks) program calculated strut lengths and angles between these vertices. These positions were also used to make a 3D mask of the clathrin structure where each strut was represented by a cylinder with 10-pixel height and 40-pixel width. The mask was then further dilated by a 50-pixel disk to create a planar mask that encompasses all the segmented vertices. This mask was used to isolate the pertinent volume within the Tikhonov filtered tomogram which was then used to create an isosurface representation for Figure 6b-c. Averaging for Figure 6f-j was performed on unmasked and unfiltered tomograms. Only vertices with three visible struts projecting from the center were included (N=626 vertices). The subtomograms included voxels within 32 nm of the vertex within the structure plane and three voxels above and below. Subtomograms were averaged along the Z dimension. The resulting images were rotated to align the strut of interest along the direction of strut A (Fig. 6f). After filtering to remove images based on length or angle, the remaining images were averaged together to attain cohort averages.

## Supplemental Figures

**Figure S1. Reproducibility of manual segmentation**. U-87 cells were segmented three times total: twice by the same person and once by a second person. (A) The measurements for percent membrane area occupation are shown. (B) The measurements for effective radius are shown. Mean and standard deviation are shown for each measurement. The mean and max difference from mean (max syst. error) for the three separate segmentations are also shown. For box plots, box is interquartile range, center line is median, center circle is mean, whiskers include all data excluding outliers.

**Figure S2. Further analysis of clathrin in eight different cell lines**. (A) The percent membrane area occupation of all clathrin is shown for all eight cell lines. (B) The percent area representation per structural category is shown for all eight cell lines. (C) Effective radius for spheres (R_sphere_) is plotted versus the effective radius for domes (R_dome_) for all cells. (D-F) Effective radius versus maximum Feret radius is shown for all cell types. (G) The percent of flat clathrin next to curved clathrin (# flat structures within 100 nm of curved clathrin)/(total # flat structures) is shown for each cell line. (H) The number of clathrin clusters, defined as groups of clathrin structures within 100 nm of each other, per area of membrane is shown. BS-C-1 cells exhibit a significant lower density than other cell types using one way ANOVA and Tukey’s test.

**Figure S3. Quantifying clathrin loss during curvature with live unroofing on a fluorescence microscope**. (A) A live HeLa cell is imaged with total internal reflection fluorescence (TIRF) microscopy during and after the unroofing process. A higher magnification is shown in (B). (C) A kymograph of this same movie shows that clathrin puncta remain and lose very little intensity after unroofing. (D) The background subtracted and normalized intensity of clathrin puncta is plotted over time after unroofing. N= 8 cells, 20 puncta per cell. Mean and standard deviation are shown.

**Figure S4. Cryotomograms**. All 18 flat clathrin structures used for tomogram data are shown. Isosurfaces are shown from the top of the clathrin structure to the left, and rotated 90° to view the structure from the side. Z-stack averages of select planes from Tikhonov-filtered tomograms are shown to the right. Some structures may have been cropped due to limits in isosurface rendering. The starred structures are clear examples where actin runs over top of the clathrin structure. Scale bars are 25 nm.

**Supplemental Movies 1-3. Cryotomograms**. Three tomograms from Figure 6a,k,m are shown as a stack of XY planes. Tomograms are filtered with Tikhonov deconvolution. Scale bar is 25 nm.

## Acknowledgements

This work utilized the NIH Multi-Institute CryoEM Facility (MICEF), the NIDDK CryoEM Facility, the NHLBI electron microscopy, and the NHLBI light microscopy cores. Specifically, we thank Huaibin Wang and Heifeng He for help in the CryoEM facilities, John Heuser for helpful discussion, and Ethan Tyler of NIH Medical Arts Department for the creation of Figure 7. Special thanks to the late Yael Mutsafi who helped inspire part of this work. J.W.T. is supported by the Intramural Research Program of the National Heart Lung and Blood Institute, National Institutes of Health.

## Author contributions

KAS, JWT designed experiments. BH, KAS, BP, MAM, AR, AS acquired platinum replica data. JWT, BH, KAS segmented and analyzed platinum replica data.GH and KAS wrote programs for analysis. BH performed spontaneous curvature experiments. JH, KAS, JJ designed cryo experiments. KAS, JJ acquired cryo data. KAS analyzed cryo data. KAS and BH did fluorescence experiments. KAS wrote and JWT edited the manuscript. All authors commented on the text.

