## Supplemental Figures S1-4 for "The structure and spontaneous curvature of clathrin lattices at the plasma membrane"

### Figure S1

Three separate segmentations of U-87 cells

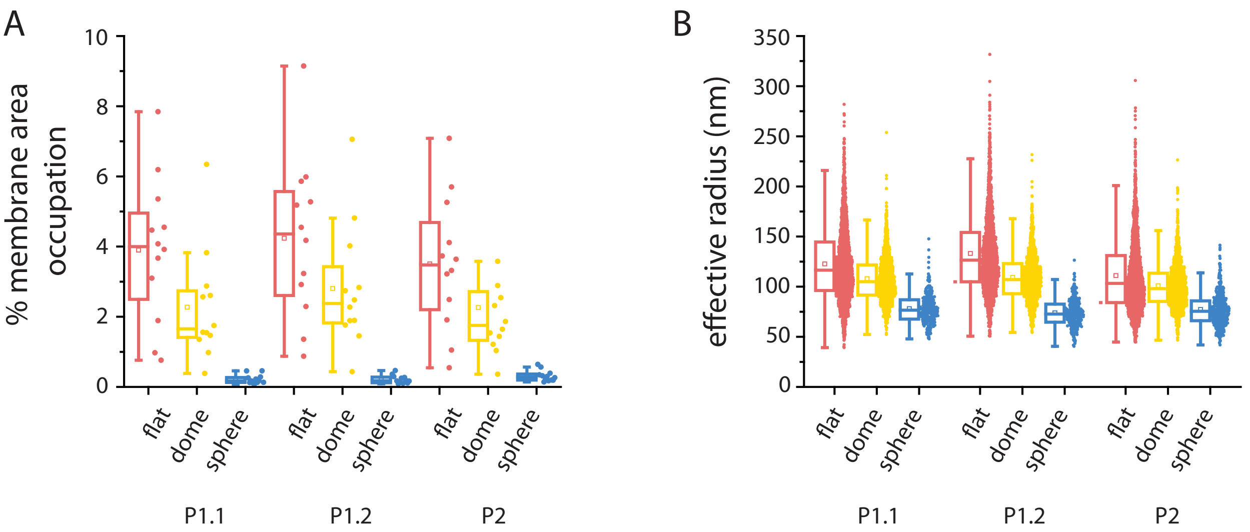


Flat % area occupation (N=12 cells)

| | | |
|---|---|---|
| P1.1 | 3.9 +/- 2.1 % | Ave. 3.9 % area<br>+/- 0.38 % max syst. error |
| P1.2 | 4.2 +/- 2.3 % | |
| P2 | 3.5 +/- 1.9 % | |

Dome % area occupation (N=12 cells)

| | | |
|---|---|---|
| P1.1 | 2.3 +/- 1.6 % | Ave. 2.4 % area<br>+/- 0.36 % max syst. error |
| P1.2 | 2.8 +/- 1.8 % | |
| P2 | 2.3 +/- 1.6 % | |

Sphere % area occupation (N=12 cells)

| | | |
|---|---|---|
| P1.1 | 0.22 +/- 0.13 % | Ave. 0.25 % area<br>0.064 % max syst. error |
| P1.2 | 0.22 +/- 0.12 % | |
| P2 | 0.31 +/- 0.16 % | |

Flat effective radius

| | | | |
|---|---|---|---|
| P1.1 | N=1996 | 122.7 +/- 37.4 nm | Ave. 122.3 nm<br>+/- 11.2 nm max syst. error |
| P1.2 | N=1732 | 133.1 +/- 40.2 nm | |
| P2 | N=2078 | 111.0 +/- 37.0 nm | |

Dome effective radius

| | | | |
|---|---|---|---|
| P1.1 | N=1329 | 107.9 +/- 23.5 nm | Ave. 106.0 nm<br>+/- 5.1 nm max syst. error |
| P1.2 | N=1574 | 109.2 +/- 23.3 nm | |
| P2 | N=1412 | 100.9 +/- 22.7 nm | |

Sphere effective radius

| | | | |
|---|---|---|---|
| P1.1 | N=300 | 78.0 +/- 14.9 nm | Ave. 76.4 nm<br>+/- 2.5 nm max syst. error |
| P1.2 | N=309 | 73.9 +/- 13.8 nm | |
| P2 | N=406 | 77.3 +/- 16.6 nm | |

### Figure S2

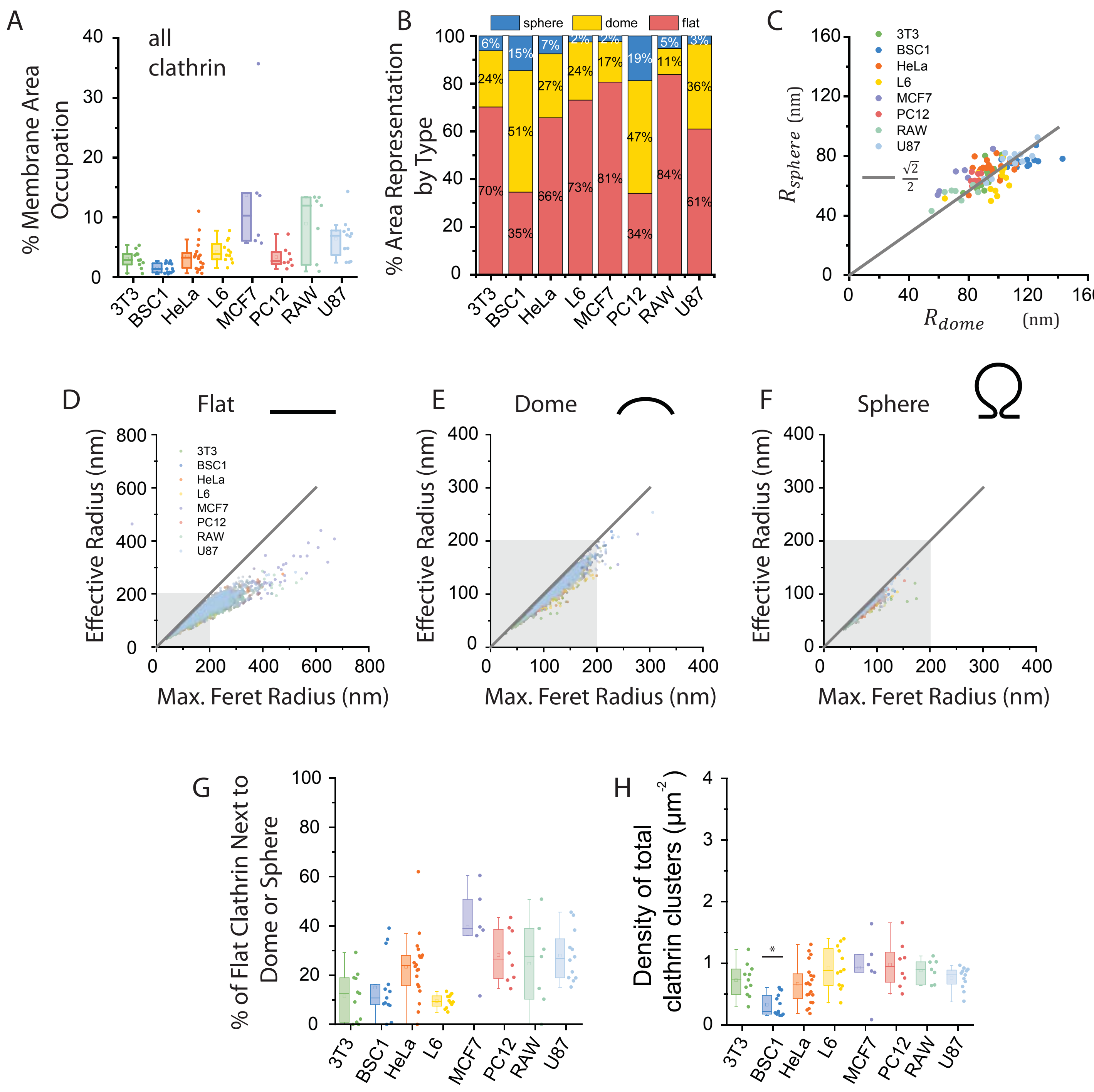
A
all
clathrin
% Membrane Area Occupation
3T3 BSC1 HeLa L6 MCF7 PC12 RAW U87
B
sphere dome flat
% Area Representation by Type
6% 24% 70%
15% 51% 35%
7% 27% 66%
2% 24% 73%
2% 17% 81%
19% 47% 34%
5% 11% 84%
3% 36% 61%
C
$R_{sphere}$ (nm)
$R_{dome}$ (nm)
$\frac{\sqrt{2}}{2}$
D
Flat
E
Dome
F
Sphere
Effective Radius (nm)
Max. Feret Radius (nm)
G
% of Flat Clathrin Next to Dome or Sphere
H
Density of total clathrin clusters (µm$^{-2}$)
*

### Figure S3

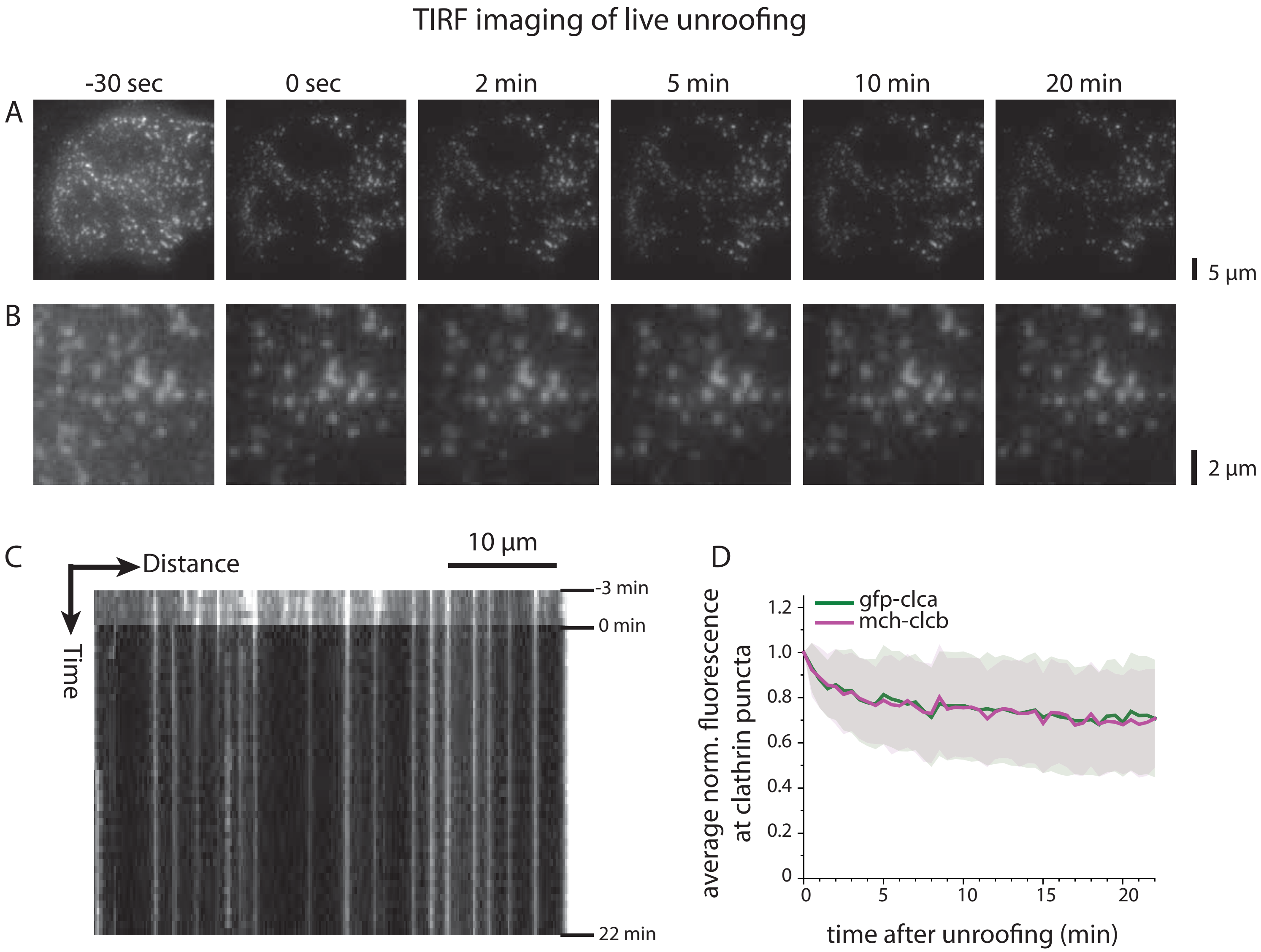

TIRF imaging of live unroofing
-30 sec
0 sec
2 min
5 min
10 min
20 min
A
5 µm
B
2 µm
C
Distance
Time
10 µm
-3 min
0 min
22 min
D
gfp-clca
mch-clcb
average norm. fluorescence at clathrin puncta
time after unroofing (min)
1.2
1.0
0.8
0.6
0.4
0.2
0
5
10
15
20

Figure S4

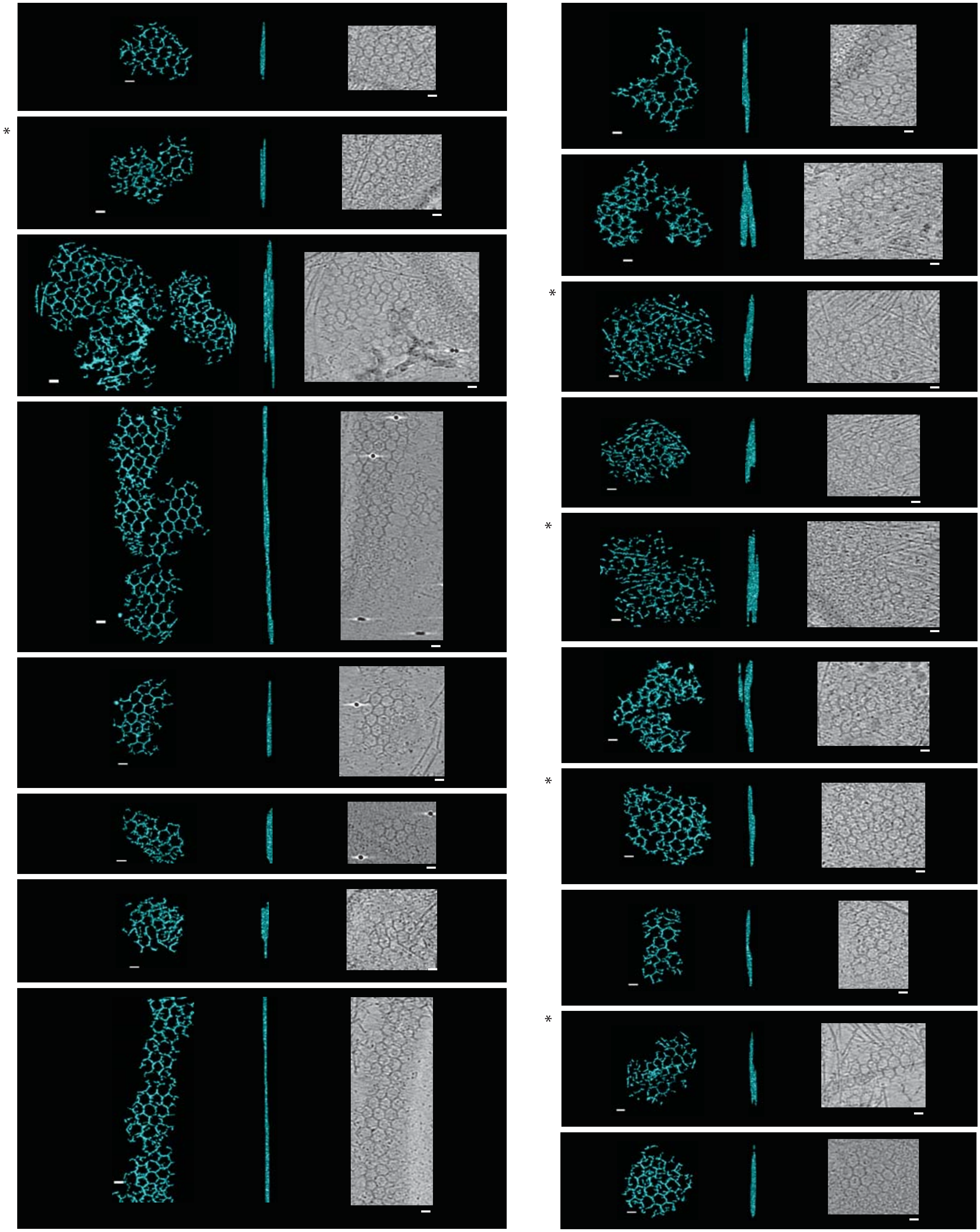
